# International and regional spread of carbapenem-resistant *Klebsiella pneumoniae* in Europe

**DOI:** 10.1101/2023.09.05.556311

**Authors:** Mabel Budia Silva, Tomislav Kostyanev, Stefany Ayala Montaño, Jose Bravo Ferrer-Acosta, Maria Garcia-Castillo, Rafael Cantón, Herman Goossens, Jesus Rodriguez-Baño, Hajo Grundmann, Sandra Reuter, the EURECA/WP1B Group

## Abstract

*Klebsiella pneumoniae,* an opportunistic pathogen able to cause hospital- and community-acquired infections, is of particular concern due to the active spread of antibiotic resistance genes associated with mobile genetic elements such as plasmids and transposons. Carbapenem-resistant *K. pneumoniae* (CRKP) exhibit a higher mortality rate than carbapenem-susceptible *K. pneumoniae* infections. In this COMBACTE-EURECA study, we present genomic data of 687 carbapenem-resistant strains recovered among clinical samples from 41 hospitals in nine Southern European countries (2016-2018) and compared these with the previous EuSCAPE collection (2013-2014). All isolates were resistant (EUCAST criteria) to at least one carbapenem antibiotic. We identified 11 clonal complexes (CCs), with most isolates belonging to the high-risk clones CC258, CC101, CC11, and CC307. *bla*_KPC-like_ was the most prevalent carbapenemase-encoding gene (45%), along with the dominance of the CC258 lineage. Equally, *bla_OXA_*_-48_ had a wide-ranging spread (39%), and *bla*_NDM_ was present in half of the ST11 isolates representing a particular lineage circulating in Greece. Two carbapenemase genes were found in 38 isolates (5.5%). Through the combination of both EURECA and EuSCAPE collections, we elucidated the evolution of *K. pneumoniae* high-risk clones circulating in Europe. Dominating clones and their associated carbapenemase genes exhibit relevant regional differences, namely CC258 in Greece, Italy, and Spain, CC101 present in Serbia and Romania and CC14 in Türkiye. Due to the wide expansion of CRKP, genomic surveillance across Europe with projects such as EURECA provides crucial insights for risk mapping at geo-temporal scales and informs necessary adaptions to the local settings for implementation of control strategies.

## INTRODUCTION

Gram-negative *Enterobacterales Klebsiella pneumoniae* (*Kp*) are part of the microbiota in humans and are considered as opportunistic pathogens able to cause hospital- and community-acquired infections such as pneumonia, bloodstream and urinary tract infections (Hansen et al., 1998). Clinically these infections are treated with fluoroquinolones, aminoglycosides, cephalosporins, and as a last resort carbapenem antibiotics. However, *Kp* may become resistant to carbapenems mainly through the acquisition of resistance genes encoding carbapenemases, or the production of extended-spectrum beta-lactamases or cephalosporinases combined with porin alterations (Bush, 2018). Carbapenemase genes are of particular concern as they can spread in association with mobile genetic elements (MGE) that are part of plasmids and transposons. These carbapenemase genes, mainly *bla*_KPC_, *bla*_NDM_, *bla*_VIM_, and *bla*_OXA-48_, are often associated with particular and successful nosocomial clones, sometimes with close relationship between a lineage and the antibiotic resistance determinants (Munoz-Price et al., 2013).

The increase of carbapenem-resistant *Kp* (CRKP) represents an unquestionable threat to the health of hospitalized patients globally with a high mortality rate compared to patients infected with carbapenem-susceptible *Kp* (Borer et al., 2009; Wu et al., 2022). Indeed, CRKP is a relevant public health problem with economic effects (Cassini et al., 2019), and it was recognized as a critical pathogen by the WHO for the need of new antimicrobial (Tacconelli, 2017). Different surveillance programs are collaboratively running at several hospitals in many countries. In Europe, national outbreaks of CRKP have been reported mainly in Southern European countries such as Greece, Spain, and Italy where also the highest prevalence is observed (David et al., 2019; Lo Fo Wong et al., 2022).

To limit the high incidence, increasing spread and unravel transmission pathways of CRKP, it is relevant to study their characteristics including distribution of lineages and resistance determinants at regional and local levels as well as internationally. Furthermore, study of such isolates may shed light on how to design better targeted antibiotic stewardship programmes for treating infections, together with the implementation of public health measures of action aiming to control further spread of CRKP (Lo Fo Wong et al., 2022).

The COMBACTE consortium pursues the prevention and treatment of antibiotic-resistant-associated infections through four main projects. Among them, COMBACTE-CARE, seeks to support the development of new treatment options, together with the analysis of clinical and epidemiological datasets in all European member states and affiliated countries (Gutiérrez-Gutiérrez et al., 2017; Kostyanev et al., 2016). As part of this, the European prospective cohort study on *Enterobacterales* showing resistance to carbapenems (EURECA) aimed to understand how the patients across Europe are infected and currently treated for *Enterobacterales*-associated infections, but also which subgroups of patients responded well to different antibiotic treatments (Gutiérrez-Gutiérrez et al., 2017). Local laboratories submitted carbapenem non-susceptible isolates to EURECA from May 2016 to November 2018 from cohorts of patients with bacterial infections in Southern European countries. Our task was to characterize and identify circulating clones of CRKP by analysing 687 genomes. In addition, we contextualized the spread of CRKP for a broader population view by comparing our data to the previous EuSCAPE study (David et al., 2019), which included the carbapenem-susceptible and non-susceptible isolates during 2013-2014 sampled from a wider range of European countries.

## MATERIALS AND METHODS

### Isolate collection and antimicrobial susceptibility

Isolates were recovered from patients diagnosed with bloodstream, intra-abdominal, pneumonia and complicated urinary tract infection, in 41 hospitals in nine countries (Albania, Croatia, Greece, Italy, Montenegro, Romania, Serbia, Spain, and Türkiye). The collection period was from May 2016 to November 2018. Overall, 49% of the samples were from blood (n=334), 31% from urine (n=213) and 20% from other sources (n=140) (Supplementary Table 1). The countries with the major sample contributions to this study were Spain (n=202), Greece (n=174), Italy (n=111), and Serbia (n=110). *Enterobacterales* isolates with MIC ≥1 mg/L (dilution methods) or ≤22 mm (disc diffusion, 10 μg disks) for meropenem or imipenem isolated from patients with the above infections were considered as putative CRE and studied; those producing carbapenemases and/or showing resistance to imipenem or meropenem according to EUCAST breakpoints were included. Collected isolates were phenotypically confirmed by disc diffusion at the EURECA central laboratory at the University of Antwerp. Further, the MIC values were determined by broth microdilution at SERMAS laboratory in the Hospital Ramon y Cajal in Madrid. The isolates were identified to the species level with MALDI-ToF, and then confirmed by whole genome sequencing.

### Whole genome sequencing and quality control

DNA extraction was performed using the Roche High Pure Template Preparation Kit. DNA concentration was measured with the Qubit fluorometer, followed by the sequencing library preparation using Nextera™ DNA Flex Library Prep (flow cell for 2x150bp paired-end sequencing).

Raw sequence reads were mapped to the reference genome MGH78578 (CP000647) using Smalt (Ponstigl, 2015) and Samtools (Danecek et al., 2021), with subsequent filtering with Genome Analysis Toolkit (McKenna et al., 2010). The minimum accepted average coverage was 30X per sample. The reads were *de novo assembled* using SPAdes v3.13.1 (Bankevich et al., 2012) with Kmer sizes 21, 33, 55, 77, 99, 109, and 123. The expected size for the assemblies was between 5-7Mb, 50-100 contigs, and N50 >100,000 bp. Kraken with minikraken database was used to check the species and potential contamination of samples (Wood & Salzberg, 2014). Sequence types (ST) were determined using the multilocus sequences typing (MLST) software (https://github.com/tseemann/mlst). The average coverage was 50x with a genome size of 5.6 Mb and N50 of 259,285 bp. The median number of contigs was 86 (Supplementary Table 1). Sequencing data has been submitted to the ENA project PRJEB63349, with individual accession identifiers for reads and assemblies given in Supplementary Table 1.

Acquired antimicrobial resistance genes were identified using Abricate v0.9.8 (https://github.com/tseemann/abricate), with a local database based on ResFinder (Zankari et al., 2012). To search mutation of *bla*_KPC_ gene in KPC producers we used BLAST (Altschul et al., 1990) after the extraction of the contig carrying the gene of interest.

### Phylogenetic analyses

Phylogenetic trees of the whole collection and subsets of particular CCs were estimated using RAxML based on SNP alignments after mapping to the reference genomes and removal of recombinant regions using Gubbins v2.4.1 (Croucher et al., 2015). Reference genomes for species overview: MGH78578 (accession number CP000647), CC11: F64 (VILG01), CC14: KPN528 (CP020856), CC15: P35 (CP053041), CC101: Kp_Goe_33208 (CP018447), CC147: HKP0064 (JACTAR01), CC258: 30660 (CP006923), CC307: CPKp1825 (WMHT01). Phylogenetic tree visualization was done with iToL (Letunic & Bork, 2021). Public datasets of interest (EuSCAPE, Italian collection of ST147 (Martin et al., 2021), Serbian collection of ST101 (Palmieri et al., 2020), worldwide collection of ST258 (Adler et al., 2017; Bowers et al., 2015; Cerqueira et al., 2017; DeLeo et al., 2014) were downloaded from the European Nucleotide Archive (ENA) and included in the analysis.

## RESULTS

### Presence of carbapenemase genes across different ST

In the EURECA collection, a total of 683 isolates were classified by the phylogenetic analysis into *Klebsiella pneumoniae* sensu stricto, 2 in *Klebsiella variicola* and 2 in *Klebsiella quasipneumoniae*. Of all the *Kp* sensu stricto we identified 50 different sequence types (STs) and 9 novel single locus variants (SLV) (Supplementary Table S1). Most isolates (n=599, 87%) were grouped in one of 11 clonal complexes (CCs; Table 1), with CC258 being the dominant lineage (n=204; 30%), and ST512 the main single ST (n=116, 17%) (Figure 1). Other CCs in order of prevalence were CC11, CC101, CC307, CC15, and CC147. Of the carbapenemase genes, *bla*_KPC-like_ (n=314; 46%) were the most abundant, mainly linked to CC258 (Figure 1). *bla*_OXA-48-like_ (n=266; 39%) was the second most widespread gene found in several STs, for instance ST15, ST101, ST11. *bla*_NDM-1_ was found in 96 (14%) isolates either on its own or in combination with *bla*_OXA-48-like_ genes, and interestingly, it was present in half of the ST11 isolates. A combination of two carbapenemase genes was found in 38 isolates (5.5%), with the majority of isolates having a combination of *bla*_OXA-48_ + *bla*_NDM-1_ (n=25, 66%) (Figure 1, Table 1). Despite re-culturing under antibiotic selection, DNA extraction, and re-sequencing, we could not find a carbapenemase in 22 isolates.

**Figure 1:**
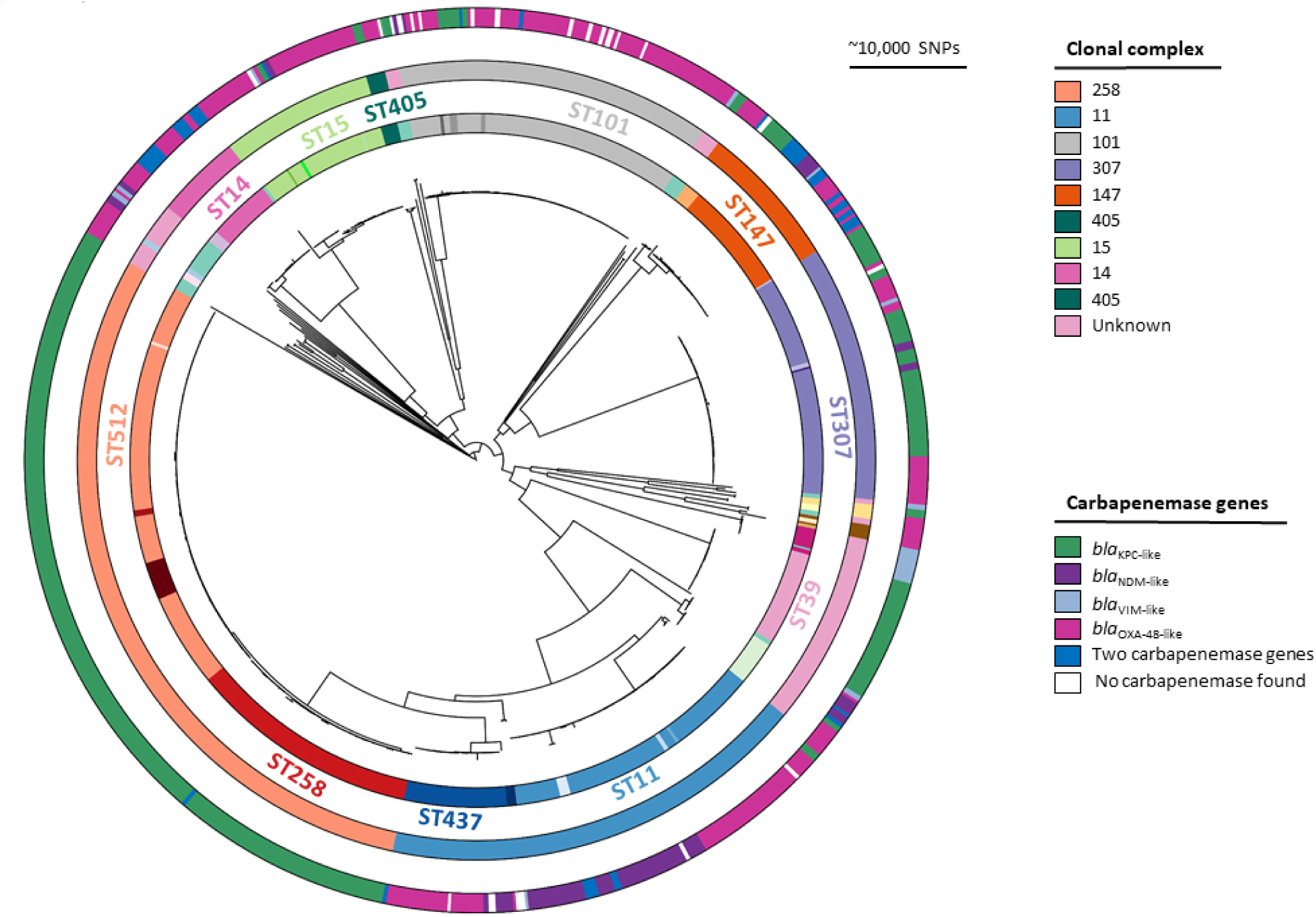
Phylogenetic overview tree of the complete CRKP EURECA collection. This phylogenetic tree contains 683 *Klebsiella pneumoniae* sensu stricto carbapenem resistant isolates collected as part of the EURECA study. The inner ring shows sequence type, the central ring displays Clonal Complex (CC), and the different colours of the outer ring depict the carbapenemase genes. Main clonal complexes with more than five isolates have been labelled.

**Table 1:** Characteristics of all submitted CRKP with different carbapenemase genes. Full data available in Supplementary Table S1.

| Clonal complex | Total number of isolates (% of total) | Countries with more than 10 isolates | STs with more than 10 isolates | Carbapenemase gene variants and number of isolates |  |  |  | No carbapenemase found |
| --- | --- | --- | --- | --- | --- | --- | --- | --- |
|  |  |  |  | Class A | Class B | Class D | Two carbapenemases |  |
| 258 | 204 (30%) | Italy (n=74)<br>Greece (n=58)<br>Spain (n=58) | ST512 (n=116)<br>ST258 (n=73)<br>ST1519 (n=12) | *bla*<sub>KPC-2</sub> (n=68)<br>*bla*<sub>KPC-3</sub> (n=135) | *bla*<sub>NDM-1</sub> (n=1) |  | 1 | 1 |
| 11 | 116 (17%) | Greece (n=41)<br>Serbia (n=37)<br>Spain (n=36) | ST11 (n=78)<br>ST437 (n=33) | *bla*<sub>KPC-2</sub> (n=4) | *bla*<sub>NDM-1</sub> (n=47)<br>*bla*<sub>VIM-1</sub> (n=1) | *bla*<sub>OXA-48</sub> (n=47)<br>*bla*<sub>OXA-48-like</sub> (n=13) | 3 | 7 |
| 101 | 87 (15%) | Serbia (n=56)<br>Romania (n=17) | ST101 (n=83) | *bla*<sub>KPC-2</sub> (n=2)<br>*bla*<sub>KPC-3</sub> (n=5) | *bla*<sub>NDM-1</sub> (n=4) | *bla*<sub>OXA-48</sub> (n=66) | 2 | 12 |
| 307 | 71 (10%) | Spain (n=33)<br>Italy (n=24) | ST307 (n=68) | *bla*<sub>KPC-2</sub> (n=16)<br>*bla*<sub>KPC-3</sub> (n=27) | *bla*<sub>NDM-1</sub> (n=4)<br>*bla*<sub>VIM-1</sub> (n=1) | *bla*<sub>OXA-48</sub> (n=22) |  | 1 |
| 15 | 42 (6%) | Spain (n=34) | ST15 (n=38) | *bla*<sub>KPC-2</sub> (n=1)<br>*bla*<sub>KPC-3</sub> (n=1) | *bla*<sub>NDM-1</sub> (n=2)<br>*bla*<sub>VIM-1</sub> (n=1) | *bla*<sub>OXA-48</sub> (n=24)<br>*bla*<sub>OXA-48-like</sub> (n=13) | 1 | 1 |
| 147 | 40 (6%) | Spain (n=20)<br>Greece (n=15) | ST147 (n=35) | *bla*<sub>KPC-2</sub> (n=12)<br>*bla*<sub>KPC-3</sub> (n=1) | *bla*<sub>NDM-1</sub> (n=13)<br>*bla*<sub>VIM-1</sub> (n=6) | *bla*<sub>OXA-48</sub> (n=18)<br>*bla*<sub>OXA-48-like</sub> (n=3) | 14 | 1 |
| 14 | 24 (4%) | Türkiye (n=24) | ST14 (n=20) |  | *bla*<sub>NDM-1</sub> (n=12) | *bla*<sub>OXA-48</sub> (n=21)<br>*bla*<sub>OXA-48-like</sub> (n=3) | 12 |  |
| 405 | 5 |  |  | *bla*<sub>KPC-2</sub> (n=2) |  | *bla*<sub>OXA-48</sub> (n=3) |  |  |
| 37 | 4 |  |  |  |  | *bla*<sub>OXA-48</sub> (n=4) |  |  |
| 45 | 4 |  |  | *bla*<sub>KPC-3</sub> (n=2) |  | *bla*<sub>OXA-48</sub> (n=2) |  |  |
| 17 | 2 |  |  |  |  | *bla*<sub>OXA-48</sub> (n=2) |  |  |
| unknown | 84 (12%) | Greece (n=47)<br>Spain (n=10)<br>Türkiye (n=10) | ST39 (n=30)<br>ST395 (n=14) | *bla*<sub>KPC-2</sub> (n=33)<br>*bla*<sub>KPC-3</sub> (n=4) | *bla*<sub>NDM-1</sub> (n=13)<br>*bla*<sub>VIM-like</sub> (n=14) | *bla*<sub>OXA-48</sub> (n=24)<br>*bla*<sub>OXA-48-like</sub> (n=1) | 5 |  |

### Geographic distribution of major clonal complexes and carbapenemase genes

We compared EURECA isolates to the CRKP isolates (n=944) of the EuSCAPE collection recovered during 2012 -2014 (David et al., 2019). The top nine CCs were the same, with minor changes in the order. Overall, the most prominent clonal complex was CC258, then CC11 and CC101, with prominent regional differences in abundance. Italy and Greece were dominated by CC258 and associated with *bla*_KPC-like_ genes (Figure 2). Greece had a higher diversity of clones with CC11-*bla*_NDM_ being prominent. In Eastern Europe, in Serbia and Romania, the abundant clone was CC101 bearing the *bla*_OXA-48_ gene, although Serbia and Romania had a second dominant clone (CC11, and CC15, respectively). Other countries, such as Spain and Türkiye displayed a more mixed picture, with several clones at near-equal proportions (Figure 2b). Common to all these countries (except Greece and Italy) is an extensive dominance of *bla*_OXA-48_ gene, which is due to the gene being carried on a promiscuous plasmid that can be present in various lineages.

**Figure 2:**
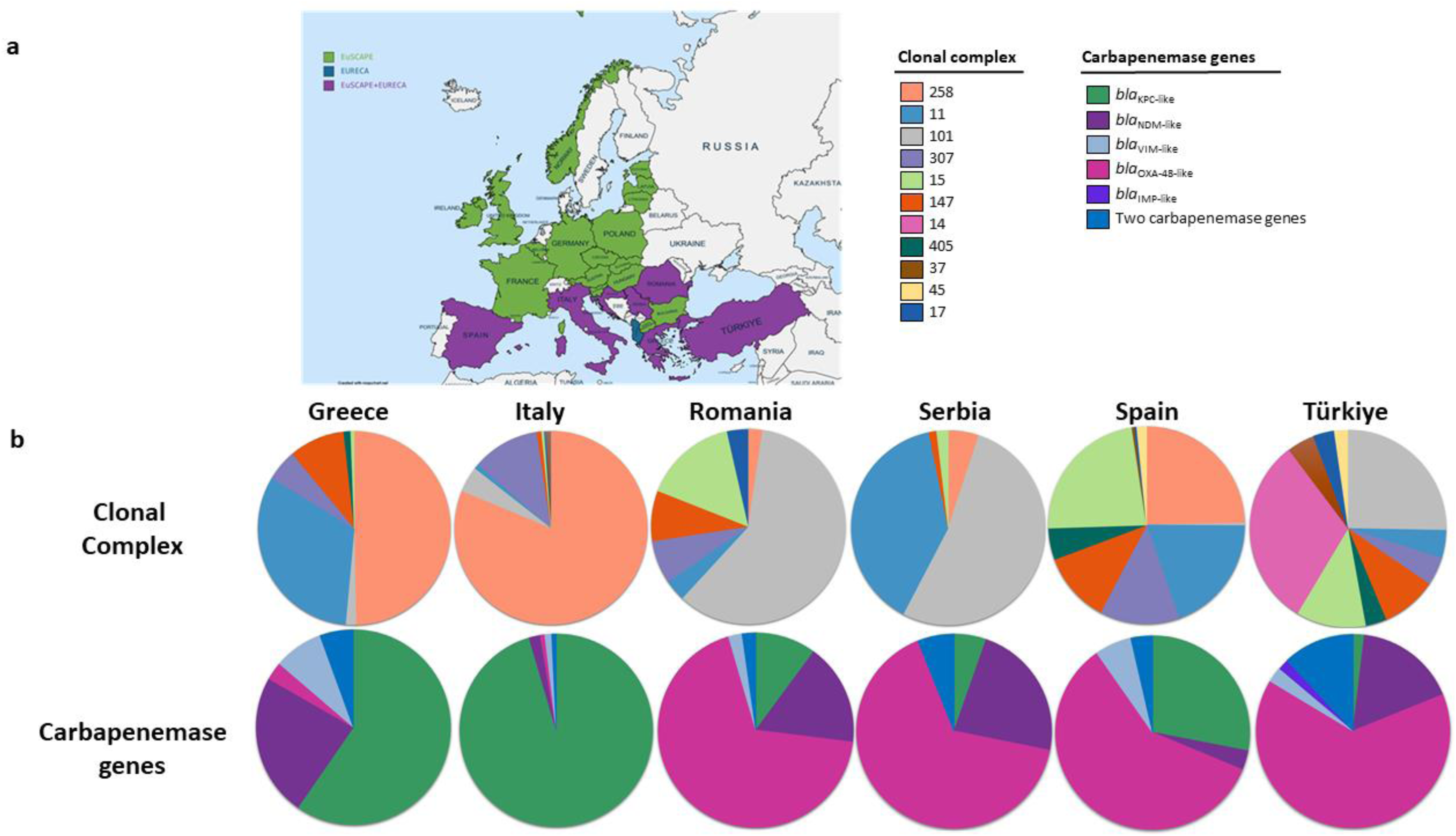
Comparison of EuSCAPE and EURECA surveys of carbapenem resistant *K.pneumoniae*. **a.** European countries, which participated in EuSCAPE (green), EURECA (blue), and the overlap of both surveys (purple). **b.** Pie charts showing the proportion of clonal complexes and carbapenemase genes of the combined EuSCAPE and EURECA surveys.

### CC258 – the worldwide spread of a successful lineage

The dissemination of the CC258 has been characterized to a large extent in previous studies. We therefore analysed the EURECA collection (Figure 3, highlighted in the first column in light blue) together with 652 publicly available genomes around the world (Adler et al., 2017; Bowers et al., 2015; Cerqueira et al., 2017; DeLeo et al., 2014), including the EuSCAPE collection (David et al., 2019). It has been reported that the ST258 first emerged in the US with *bla*_KPC-2_ (light green isolates). A subsequent introduction in Europe started in Greece with *bla*_KPC-2_, from where it expanded towards Serbia and Montenegro. An independent introduction occurred in Romania. Then, ST258 acquired the *bla*_KPC-3_ gene, and there was a small introduction into Italy visible in the Bowers *et al* 2015 and EuSCAPE datasets but not within EURECA. Another introduction of ST258 with *bla*_KPC-3_ gene happened in Israel. A second introduction from there into Greece took place, showing fewer isolates than the lineage with the *bla*_KPC-2_ gene. In Israel, the lineage evolved into ST512, this being the lineage that subsequently spread successfully from there onwards, particularly in Italy (Figure 3). Some single locus variants (SLV) of ST512 have arisen there, ST868 (red) and ST1519 (orange). From Italy, a clear introduction is now visible into Spain, which was lacking in the EuSCAPE dataset. This may be due to sampling in a different city than before (Figure 3).

**Figure 3:**
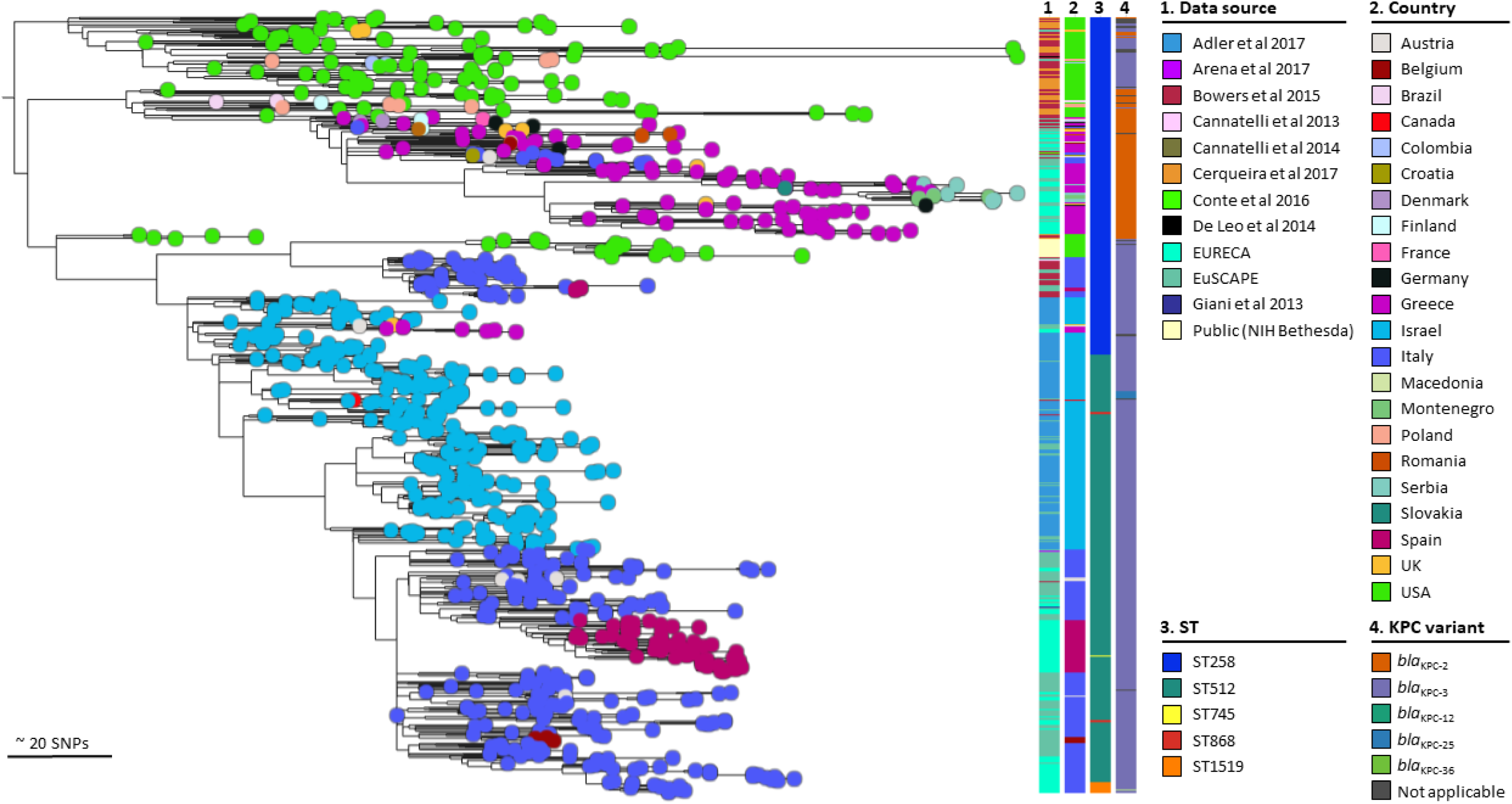
Global spread of the epidemic CC258. A phylogenetic tree of 856 isolates of ST258, ST512 and single locus variants with 204 isolates from EURECA, 236 isolates submitted to EuSCAPE, and 415 isolates with publicly available sequence data. Columns indicate: Project (1), country (2), ST (3), and KPC-variant (4). For further metadata and interactive exploration, follow this Microreact link: https://microreact.org/project/cxmkXctuhnWx4vAbCbtTFF-global-st258512-klebsiella-pneumoniae

Looking for the new mutations of *bla*_KPC_ gene, we found only one isolate (1/12) of ST1519 carrying *bla*_KPC-36_ gene that confer resistance to cefatzidime/avibactam (Figure 3).

### ST307 – the emergence of a new threat?

ST307 is an emerging clone with a potential of adapting into a successful lineage (Villa et al., 2017; Wyres et al., 2019). For this reason, we specifically analysed the phylogenetic relationship of 117 isolates CC307 (ST307 and 3 different SLV) from the EuSCAPE and EURECA collections. The highest circulation of this clone was in Greece, Italy, and Spain (Table 1). Isolates from Spain grouped in different clades compared to other countries like Greece, Italy, and Romania (Figure 4a). We investigated the circulating clones in detail in these countries by reconstructing two different phylogenetic trees, and used the cutoff of 21 SNP value previously proposed for the discrimination of hospital clusters (David et al., 2019). Isolates with less than 21 SNPs were indeed isolated from the same hospitals, and we confirmed larger SNP distances between different countries, indicative of local circulating clones rather than direct clonal transmissions (Figure 4b and c). The most prevalent carbapenemase gene was *bla*_KPC-like_, particularly in Italy and Greece, which also corresponded to the most prevalent carbapenemase type in these countries (Figure 2b, 4b). Previous studies have also shown the association of CRKP ST307 with the *bla*_KPC_ gene (Loconsole et al., 2020). In Spain, small outbreaks within single hospitals were observed (Figure 4c). Here, most isolates carried the *bla*_OXA-48_ gene, or a combination of *bla*_KPC-3_ and *bla*_OXA-162_ (Figure 4c).

**Figure 4:**
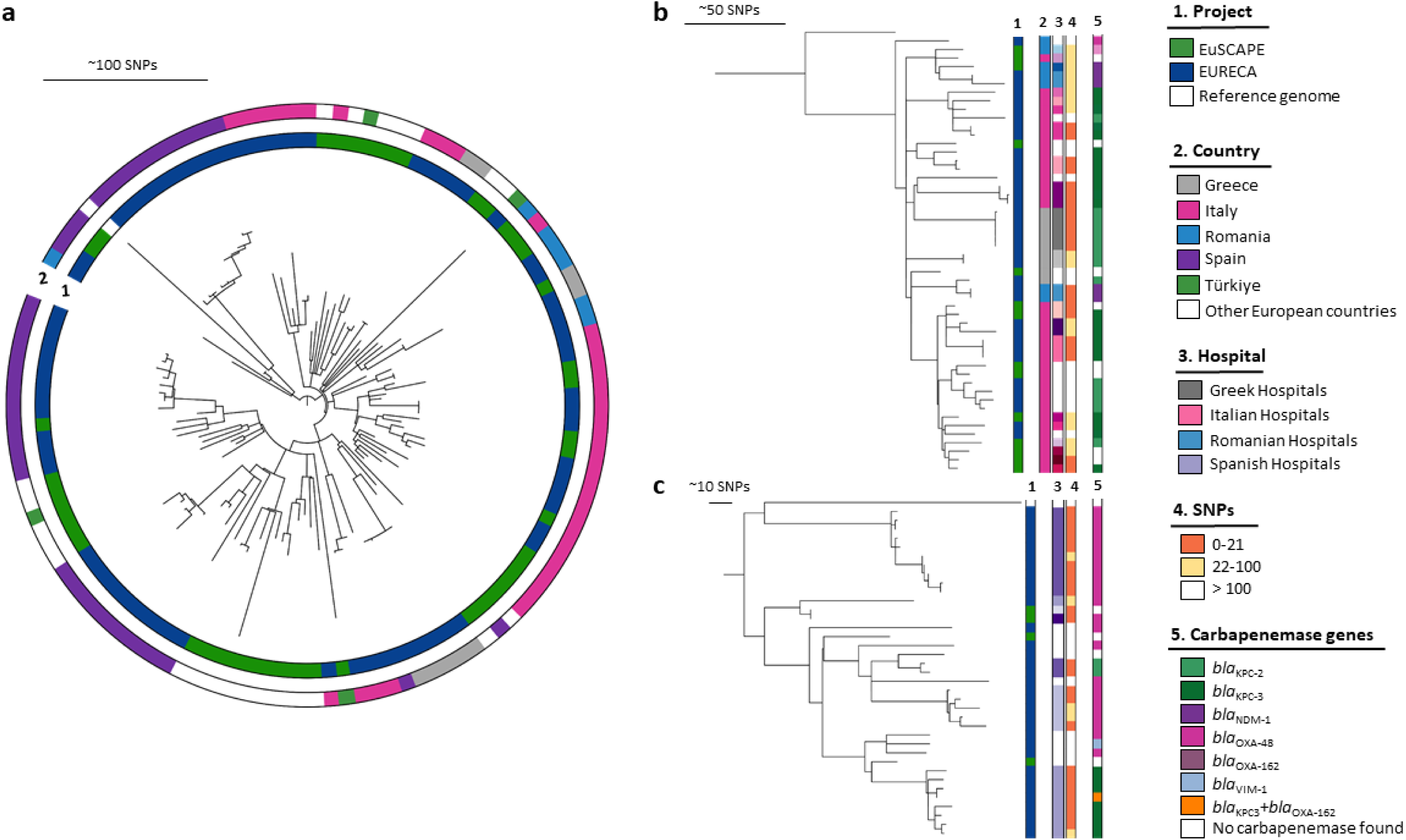
Phylogenetic analysis of CC307. **a.** Combined analysis of 117 isolates from EuSCAPE and EURECA. Inner circle depicts the study, outer circle country of origin **b.** Phylogenetic relationship between isolates CC307 from Greece, Romania and Italy, **c.** Isolates CC307 from Spain. Columns present the information about: 1 – Project, 2 – Country 3 – Different hospitals (shades for each country) 4 – SNPs distance 5 – Carbapenamase genes.

### CC11 and CC101 – the importance of knowing local conditions

A total of 330 isolates of the CC11 from the EuSCAPE and EURECA collections were analysed. The phylogenetic tree formed three clades (Figure 5a): the first clade containing the isolates belonging exclusively to ST437, a second clade with ST11 isolates plus a nested ST340 clade, and a third clade with ST11 isolates only. The distribution of carbapenemase genes differed between the countries, for instance in Greece the clone ST11 was associated with the *bla*_NDM-1_ gene. In contrast, in Spain *bla*_OXA-48_ as well as *bla*_OXA48-like_ genes were present (Figure 5a). Overall, the isolates clustered largely by country and no transmission events were observed between countries.

**Figure 5:**
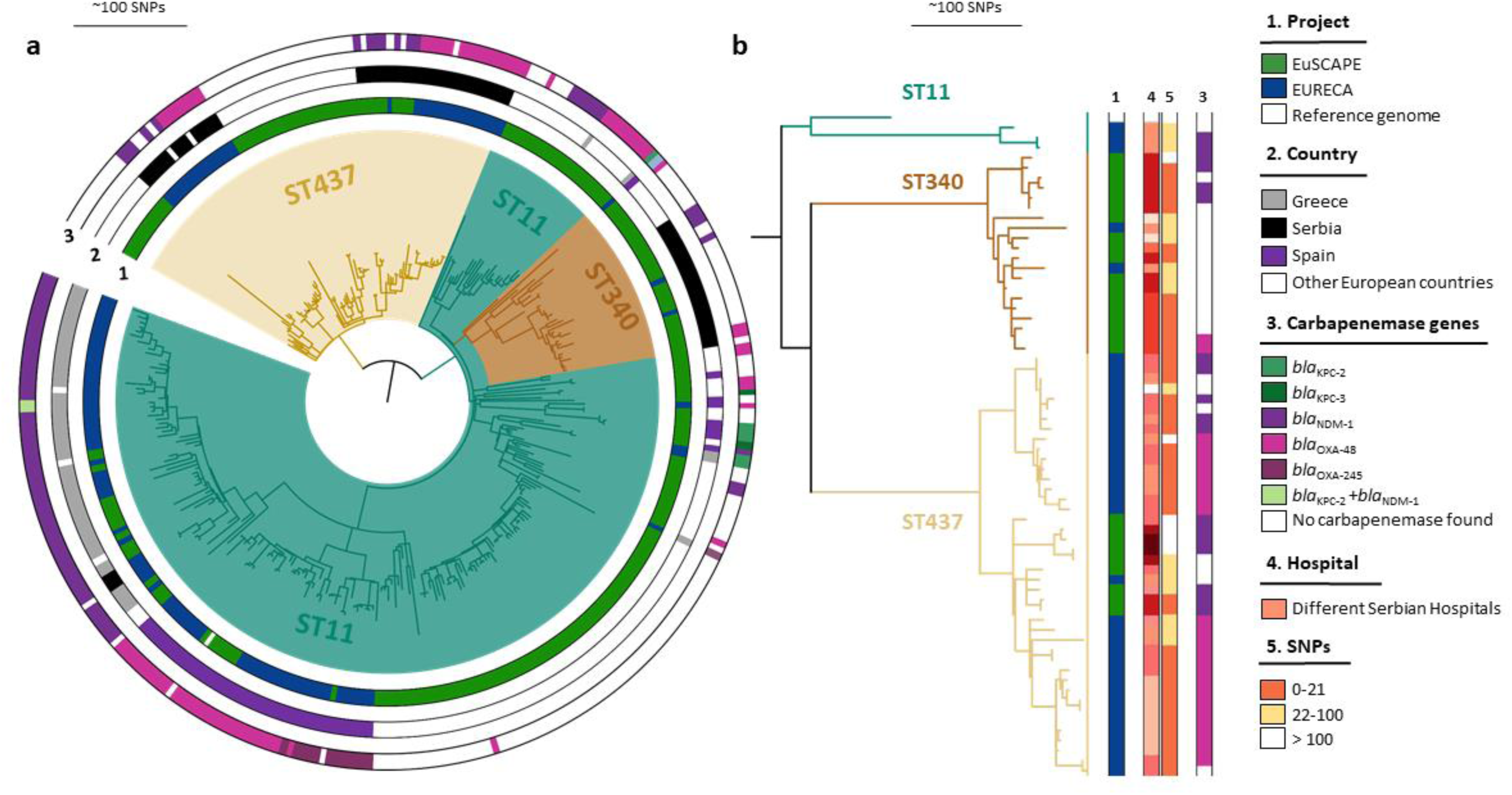
Phylogenetic analysis of CC11. **a.** Combined analysis of 329 isolates from EuSCAPE and EURECA. Inner ring: project, middle ring: geographic origin, outer ring: carbapememase genes. **b.** Phylogenetic relationship between isolates CC11 from Serbia only. Columns display the information about project (1), different colours representing different hospitals in Serbia (4), SNP distance (5), and carbapenemase genes (3).

For an in-depth study, we reconstructed a phylogenetic tree only with the Serbian isolates, due to most of them belonging to two different STs to the founder ST11 (Figure 5b). Interestingly, the ST340 clade contained predominantly carbapenem-non-susceptible EuSCAPE isolates without carbapenemase genes but with extended-spectrum betalactamase *bla*_CTX-M-15_. In this clade, both highly clonal isolates within the same hospital (0-21 SNPs), and signs of a nationally spreading clone (22-100 SNPs for isolates collected from different hospitals) are visible. Isolates within the ST437 clade frequently carried either *bla*_NDM-1_ or *bla*_OXA-48_. Two of the subclades belonged exclusively to the EURECA collection and might represent a better approximation of the current circulating clones of carbapenem-non-susceptible ST437 in Serbia (Figure 5b).

In CC101 (Supplementary Figure 1a), EURECA isolates mainly came from Serbia and Romania and formed separate clades. The Romanian isolates appeared to be more diverse and were in a clade with isolates from Italy, Spain and Türkiye. In contrast, isolates from Serbia formed distinct clades, so we investigated them in more detail. To define the ancestral relationships of the CC101 in Serbia, we analyzed 135 genomes: 57 of EURECA collection, 42 from the EuSCAPE survey (David et al., 2019), and 36 genomes from Palmieri et al. (Palmieri et al., 2020). The latter study recovered colistin resistance CRKP of ST101 between 2013 and 2017 and confirmed MgrB mutations as the major cause of colistin resistance in these isolates. They also reported *bla*_OXA-48_ as the carbapenemase gene endemic in Serbia, which agrees with our results. The phylogenetic tree showed several clades, with the EuSCAPE isolates being the oldest and forming the most basal clade. These isolates were also largely carbapenem-susceptible and did not harbour a carbapenemase (with one exception) (Supplementary Figure 1b). There are further, more clonal EuSCAPE isolates with either *bla*_OXA-48_, *bla*_NDM-1_ or a combination of both, which form the basis for a clonal expansion seen in the EURECA and Palmieri collections. This clonal expansion (less than 21 SNPs between isolates) was characterized by the presence of *bla*_OXA48_, and might be indicative of local transmission chains. However, we cannot confirm this since we have no further information about the isolates. Importantly, we could rule out that these isolates contributed twice to the different collections because the time frames did not overlap.

### Other clonal complexes

Regarding other CCs, one to highlight is the **CC147**, which was mainly recovered from Spain and Greece in the EURECA collection. An outbreak in Tuscany, Italy 2018 (Martin et al., 2021), had been reported and therefore we contextualized our isolates plus the EuSCAPE isolates (Supplementary Figure 2). EuSCAPE and EURECA isolates were mostly diverse, with only two small clusters in Greece and Serbia (EURECA collection) indicative of local clones, whereas the Tuscany outbreak formed a separate, highly clonal expansion. Interestingly, the carbapenemase genes present showed a wide variety even in the local lineages, only the Tuscany outbreak was very homogenous.

The **CC15** was also diverse, recovered from a wide range of countries, and with different carbapenemase combinations (Supplementary Figure 3). In Spain, we identified two clades; in the first clade isolates carried *bla*_OXA-48_ and the other, highly clonal clade with 14 isolates from the same hospital exhibited *bla*_OXA245_ exclusively (EURECA collection).

Lastly, the **CC14** from the EURECA collection highlighted a local expansion with ST14 and the single variant ST2096 in Türkiye. Most of the isolates were from the same hospital except two, carrying the *bla*_OXA-232_ and *bla*_OXA-48_ gene on its own or in combination with *bla*_NDM-1_ gene (Supplementary Figure 4).

## DISCUSSION

Due to their rapid and efficient spread, CRKP are considered a public health problem worldwide. In order to monitor the clonal evolution and geographical expansion over time, new tools such as WGS have been implemented. The characterization of carbapenem-resistant isolates is essential for infection control purposes because of their impact on therapeutic decisions in clinics, hence laboratory detection providing high accuracy and fast diagnosis is key for accurate antibiotic delivery (Hernández-García et al., 2022). The entire collection of 683 isolates in this present study (COMBACTE-CARE EURECA) exhibited phenotypic resistance to at least one carbapenem commonly used to treat infections caused by multiple drug-resistant pathogens. The collection was split in 11 clonal complexes, the most prominent being CC258, CC11, CC101, and CC307 as the major high-risk clones distributed worldwide (Figure 2a, b). As the EuSCAPE collection temporally preceded EURECA by two years, we contextualized both collections to better illustrate the clonal distribution of CRKP in Europe.

Since the first report in 2001 of the clonal lineage CC258 in the USA associated with *bla*_KPC_, its international spread has been rapid and extensive (David et al., 2019). Using isolates belonging to CC258 of our current collection along with previously published ones (Figure 3), we confirmed the continental spill-over of CC258 into Europe from the USA, which began with several introductions into Greece and Italy, the expansion into Serbia and Montenegro, Romania and finally into Spain. EURECA helped to highlight more clearly the second introduction of this clone into Spain and Serbia (Figure 3). We confirmed that this lineage linked to *bla*_KPC-like_ is still prevalent in Italy, Greece and Spain (Figure 2). One isolate of the single locus variant ST1519 harboured *bla*_KPC-36_ gene and was recovered from a blood of a hospitalized patient in Bologna-Italy, a previous study reported this novel *bla*_KPC-3_ variant in the same city (Gaibani et al., 2020). This new mutation confer resistance to ceftazidime/avibactam use for the treatment of CRKP (Haidar et al., 2017).

CC258 was not the most abundant (Bonnin et al., 2020; Rodrigues et al., 2016) in other European countries such as France and Portugal in 2014 and 2018, respectively, this could be due to the rise of other high-risk clones like CC307, CC101 and CC11.

The emerging high-risk clone ST307 was first described in the mid-1990s, and it has since been reported worldwide with different carbapenemase resistance determinants (Castanheira et al., 2013; Ma et al., 2013; Heiden et al., 2020). This clone has also been responsible for nosocomial outbreaks in European countries such as Portugal, Spain, France, UK, Germany, Netherlands, Slovenia and Romania (Peirano et al., 2020). In addition, it is considered a replacement of other high-risk clones, such as ST258 in countries like Colombia (Ocampo et al., 2015), South Africa (Lowe et al., 2019), and Italy (Bonura et al., 2015). Furthermore, it was also responsible for nosocomial outbreaks harbouring *bla*_KPC_ variants conferring ceftazidime-avibactam resistance (Hernández-García et al., 2022) and more recently cefiderocol resistance(Castillo-Polo et al., 2023). The presence of several chromosomally and plasmid-encoded factors associated with hypervirulence make ST307 a superior clone that can easily share large plasmids with other *Kp* and *Enterobacterales* species such as the *Enterobacter cloacae* complex (Heiden et al., 2020). Indeed, the presence of a distinct capsule structure (rare in the *Kp* population) may contribute to the successful propagation of this lineage (Wyres et al., 2020). In the EuSCAPE and EURECA collections we observed that there was a regional circulation of this clone in Greece, Italy and Spain (Figure 4). In our two observed collections, the carbapenemase gene found in this clone often appeared to reflect the dominant carbapenemase in that country, namely *bla*_KPC-2_, and *bla*_KPC-3_ in Greece and Italy, *bla*_OXA-48_ for the Spanish isolates. This may be a hint that the adaptability of ST307 with respect to carbapenemase uptake could be influenced by geographic prevalence, although interestingly, the Romanian ST307 isolates carried *bla*_NDM-1_ (Figure 4).

Large surveillance and sequencing efforts such as the current study not only improve our understanding of global clones, but may also highlight regional circumstances that need to be taken into account in local infection control efforts. In Serbia, two different STs of CC11 circulate: ST340, largely non-susceptible to carbapenems but without carbapenemases from the EuSCAPE collection, and ST437 mainly from the later EURECA collection, whose isolates often harbour either *bla*_OXA-48_ or *bla*_NDM-1_ (Figure 5b). The founder ST11 is of minor importance as we found few isolates belonging to this ST, and putatively only signifies a single small introduction event from Greece (Figure 5a). Comparing all non-susceptible isolates in EuSCAPE and EURECA from Serbia, the percentage of ST101 isolates has stayed the same, however the ST437 lineage has doubled in proportion. Since EURECA sampled less but completely overlapping hospitals with EuSCAPE, this may be an emerging trend amongst Serbian isolates. Also in Serbia, a particular ST101 clone appears to have emerged, that is characterized by the presence of *bla*_OXA-48_ (Supplementary Figure 1a, b). From the current collections, we are not able to say whether the susceptible clones are still circulating in Serbia, or whether ST101 isolates would now exclusively be part of the non-susceptible clade. Similarly, in Spain several different lineages of CC15 co-circulated; two lineages where isolates carried *bla*_OXA-48_ and one clade with *bla*_OXA-245_ (Supplementary Figure 3).

## CONCLUSIONS

Regional differences exist in terms of the local dominant CRKP clones. In Greece, Italy, and Spain, CC258 was the dominant clone, while, in Serbia and Romania, CC101 was predominant, and CC14 was potentially expanding in Türkiye. The respective carbapenemase genes within these CCs are likewise diverse. Overall, *bla*_KPC-like_ was the most prevalent carbapenemase gene (46%) associated with the most abundant CC258. The second most frequent carbapenemase gene was *bla*_OXA-48_ (39%) widely spread between different STs. Moreover, a relatively high proportion of isolates (5.5%) harboured two carbapenemases.

The combination of both EURECA and EuSCAPE collections helped to elucidate the high-risk clones circulating in Europe and the evolution of *Kp* through the shift of dominant lineages, their associated carbapenem resistance genes, and the potential acquisitions of locally circulating genes. Collections such as these demonstrate that continuous surveys are necessary and the sampling should always include susceptible background populations as well as highly resistant isolates in order to discern introduction from *de-novo* emergence.

These surveys also provide a blueprint for important European initiatives towards better planning for public health and infections control interventions.

## Acknowledgements

The authors wish to thank the microbiologists from all participating laboratories in the EURECA study, and the members of the EURECA/WP1B group for their dedicated work and substantial contribution during the isolate collection phase of this study. The EURECA/WP1B group members are: Begoña Palop-Borrás, Irene Gracia Ahufinger, Fe Tubau, Fernando Chaves, Julio Garcia, Rafael Cantón, Francisca Portero, Patricia Muñoz, Silva Tafaj, Arjana Tambic Andrasevic, Sophia Vourli, Efthymia Protonotariou, Eleni Vagdatli, Ioanna Voulgaridi, Iris Spiliopoulou, Ioannis Deliolanis, Maria Panopoulou, Ergina Malli, Efthymia Petinaki, Efstathia Perivolioti, Theodora Biniari, Nicholas Legakis, Antonino Di Caro, Sara Giordana Rimoldi, GianMaria Rossolini, Maria Paola Landini, Teresa Spanu, Milena Arghittù, Rossana Cavallo, Anna Marchese, Susanna Cuccurullo, Arsim Kurti, Milena Lopicic, Olivia Dorneanu, Mirela Flonta, Alma Kosa, Daniela Talapan, Mariana Buzea, Camelia Ghita, Edit Székely, Snezana Jovanovic, Teodora Vitorovic, Deana Medic, Branislava Kocic, Ana Perucica, Cüneyt Özakin, Banu Sancak, Can Bicmen, Zeynep Ceren Karahan, Ufuk Hasdemir, Guner Soyletir, Levantia Zachariadou, Anastassios Doudoulakakis, Paola Bernaschi, Roberto Bandettini and Vassiliki Pitiriga.

## Funding

TK, JBF-A, MG-C, RC, HGo, JR-B, HGr and the research leading to these results received support from the Innovative Medicines Initiative Joint Undertaking (https://www.imi.europa.eu/) under Grant Agreement Nos. 115523 [COMBACTE-NET (Combatting Bacterial Resistance in Europe)] and 115620 (COMBACTE-CARE), resources of which are composed of financial contributions from the European Union’s Seventh Framework Programme (FP7/2007-2013) and EFPIA companies’ in kind contribution. JR-B and RC receive overarching funding for research by Plan Nacional de I+D+I, Instituto de Salud Carlos III, Subdirección General de Redes y Centros de Investigación Cooperativa, Ministerio de Ciencia, Innovación y Universidades, Spanish Network for Research in Infectious Diseases (REIPI RD16/0016/0001 and RD16/0016/0011) and CIBERINFEC (21/13/00012, 21/13/00084) co-financed by the European Development Regional Fund ‘A way to achieve Europe’, Operative Program Intelligence Growth 2014–2020. MBS is supported by the JPI-AMR call (K-STaR 01KI1910), and SAM is supported by the Federal Ministry of Education and Research (BMBF; TAPIR 01KI2018). SR is a Junior Research Group Leader supported by the BMBF (TAPIR 01KI2018). We thank Valentina Valenzuela Dallos and Sarah Klassen for excellent technical support.

## Figures

**Supplementary Figure 1:**
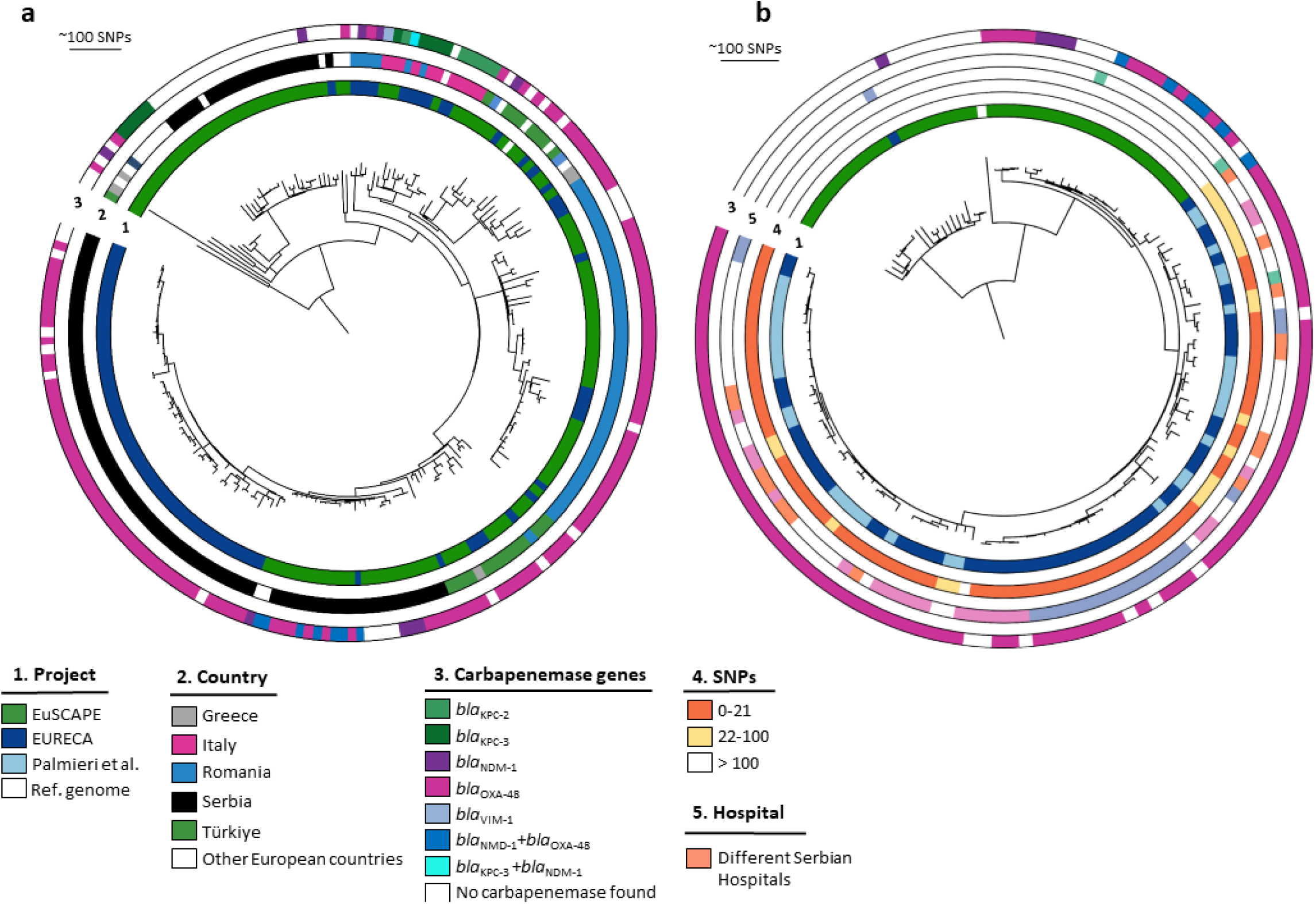
Phylogenetic analysis of CC101. **a.** Phylogenetic tree of 213 isolates from EURECA and EuSCAPE survey. Inner ring: project, middle ring: geographic origin, outer ring: carbapememase genes. **b.** 135 isolates ST101 from Serbia recovered during EuSCAPE, EURECA survey and Palmieri et al study (Palmieri et al., 2020). The rings show from inside to outside the project (1), number of SNP differences (4), different colours representing different hospitals in Serbia (5), and carbapenemase genes (3).

**Supplementary Figure 2:**
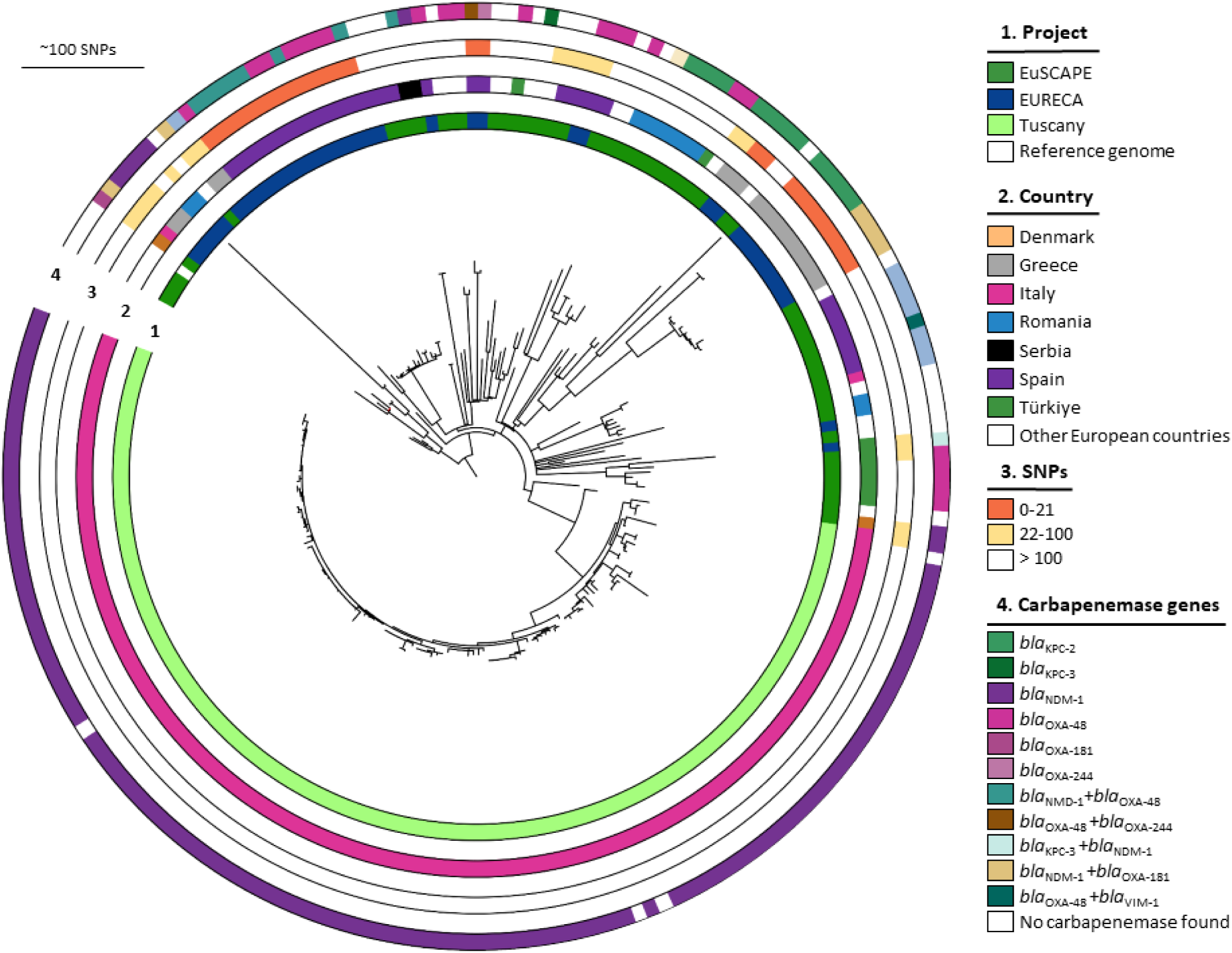
Phylogenetic analysis of CC147. Phylogenetic tree of 212 isolates from EURECA, EuSCAPE survey and an outbreak in Tuscany Italy (Martin et al., 2021). The rings show the project (1), geographic origin (2), SNP distance (3), and carbapenemase genes (4).

**Supplementary Figure 3:**
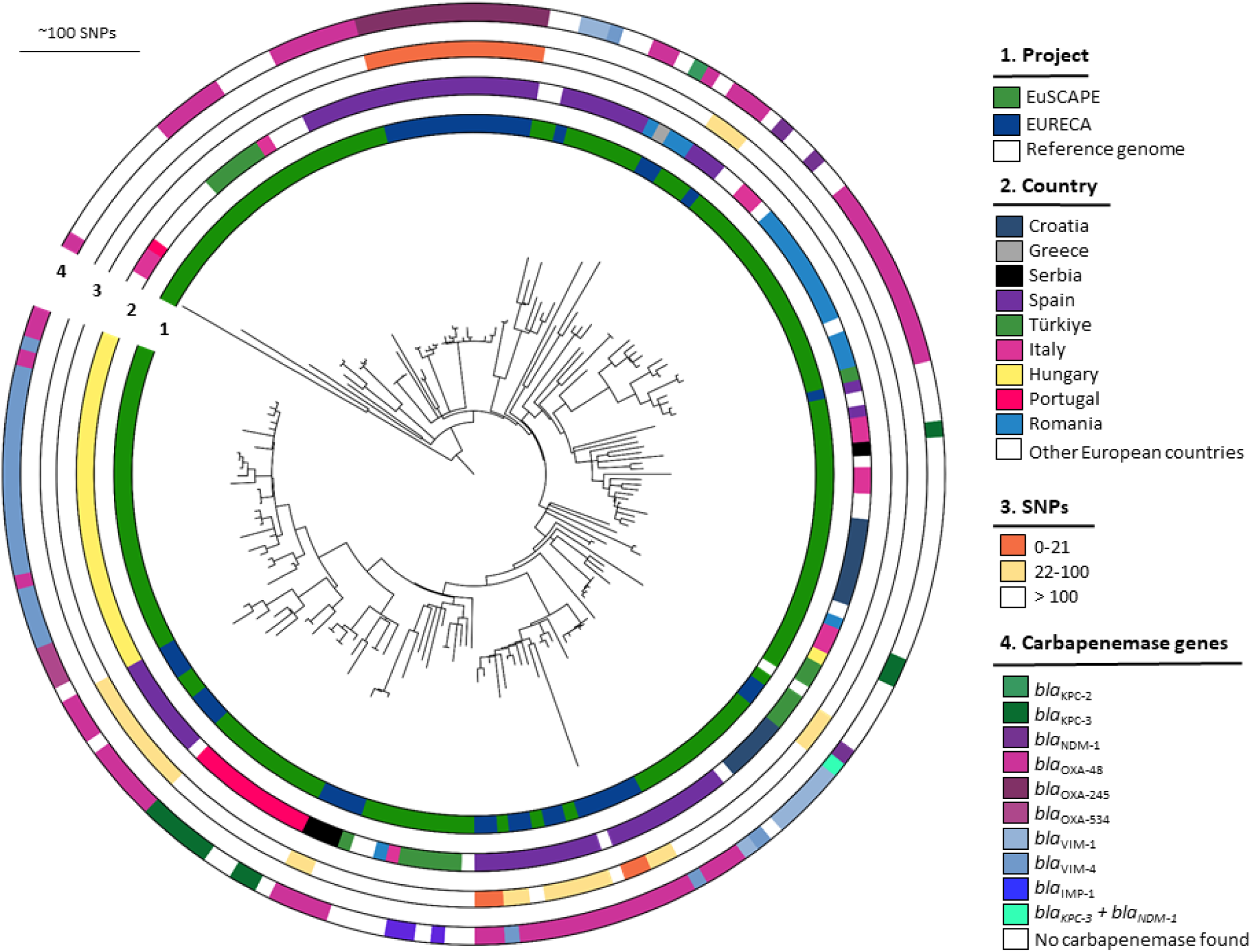
Phylogenetic analysis of CC15. Phylogenetic reconstruction of 189 isolates from EURECA and EuSCAPE survey. The rings show the project (1), geographic origin (2), SNP distance (3), and carbapenemase genes (4).

**Supplementary Figure 4:**
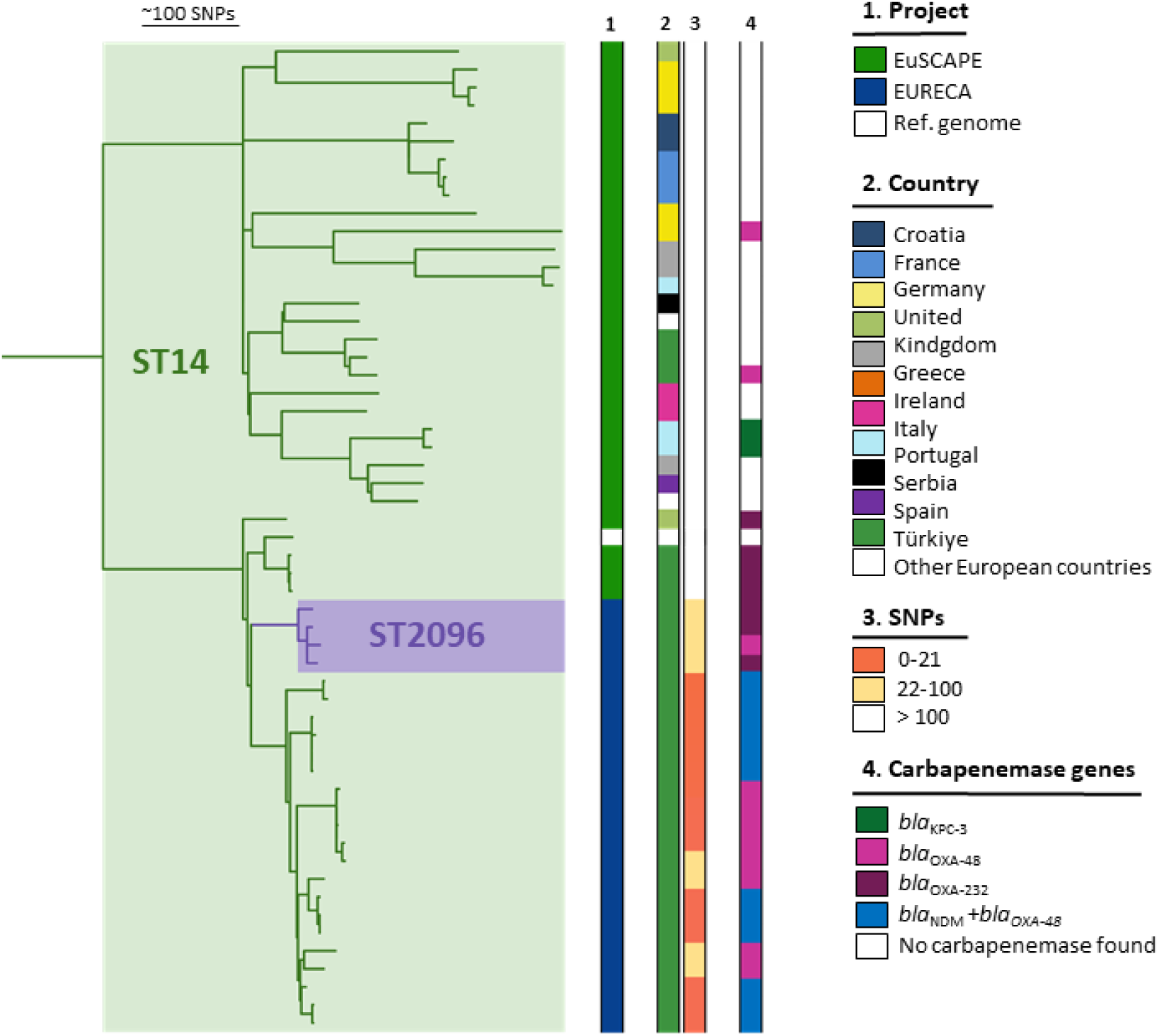
Phylogenetic analysis of CC14. Phylogenetic tree of 54 isolates from EURECA and EuSCAPE survey. The rings show the project (1), geographic origin (2), SNP distance (3), and carbapenemase genes (4).

## Notes

https://www.ebi.ac.uk/ena/browser/view/PRJEB63349

### Competing Interest Statement

The authors have declared no competing interest.

